# *Prevotella timonensis* degrades the vaginal epithelial glycocalyx through high fucosidase and sialidase activities

**DOI:** 10.1101/2024.01.09.574844

**Authors:** Celia Segui-Perez, Rivka de Jongh, Robin Jonkergouw, Paula Pelayo, Emily P. Balskus, Aldert Zomer, Karin Strijbis

## Abstract

Bacterial vaginosis (BV) is a polymicrobial infection of the female reproductive tract (FRT). BV is characterized by replacement of health-associated *Lactobacillus* species by diverse anaerobic bacteria, including the well-known *Gardnerella vaginalis*. *Prevotella timonensis* and *Prevotella bivia* are anaerobes that are found in a significant percentage of BV patients, but their pathogenic properties are yet to be determined. Defining characteristics of anaerobic overgrowth in BV are adherence to the mucosal surface and the increased activity of mucin-degrading enzymes such as sialidases in vaginal secretions. Here, we demonstrate that *P. timonensis* but not *P. bivia* strongly adhered to vaginal and endocervical cells to a similar level as *G. vaginalis* but did not elicit a comparable pro-inflammatory response. The *P. timonensis* genome uniquely encodes a large set of mucus-degrading enzymes including 4 putative fucosidases and 2 putative sialidases, PtNanH1 and PtNanH2. Enzyme assays demonstrated that fucosidase and sialidase activity in *P. timonensis* cell-bound and secreted fractions was significantly higher than for other vaginal anaerobes. Infection assays revealed that *P. timonensis* fucosidases and sialidases efficiently removed fucose and α2,3- and α2,6-linked sialic acid moieties from the epithelial glycocalyx. Recombinantly expressed *P. timonensis* NanH1 and NanH2 efficiently removed α2,3 and α2,6-linked sialic acids from the epithelial surface and sialic acid removal by *P. timonensis* could be blocked using inhibitors. This study demonstrates that *P*. *timonensis* has distinct virulence properties that include initial adhesion and a high capacity for mucin degradation at the vaginal epithelial mucosal surface. Our results underline the importance of understanding the role of different anaerobic bacteria in BV.

**Significance statement (Layman):** Bacterial vaginosis (BV) is a common vaginal infection that affects a high percentage of women and is associated with reduced fertility and increased risk of secondary infections. *Gardnerella vaginalis* is the most well-known BV-associated bacterium, but *Prevotella* species including *P. timonensis* and *P. bivia* may also play an important role. We showed that, similar to *G. vaginalis*, *P. timonensis* adhered well to the vaginal epithelium, suggesting that both bacteria could be important in the first stage of infection. Compared to the other bacteria, *P. timonensis* was unique in efficiently removing the protective mucin sugars that cover the vaginal epithelium. These results underscore that vaginal bacteria play different roles in the initiation and development of BV.

## Introduction

Bacterial vaginosis (BV) is a complex polymicrobial vaginal infection that is prevalent in women of different ages. BV is associated with increased susceptibility to sexually transmitted infections (STIs) including Human Immunodeficiency Virus (HIV) (*1*, *2*) and Human Papilloma Virus (HPV) (*3*) but also infertility (*4*), and adverse pregnancy outcomes including pre-term birth (*5*). BV is diagnosed according to the Amsel criteria that include a high vaginal pH (>4.5), detection of thin discharge, an odor of amines after addition of potassium hydroxide, and the presence of “clue cells” in vaginal secretions (*6*, *7*). Bacterial gram staining followed by the Nugent score test is also used to diagnose BV (*8*, *9*). Vaginal secretions of BV patients contain enzymes that are capable of degrading the protective mucus layer including mucinases and sialidases that can also be used for diagnostics (*10–12*).

In contrast to BV, health-associated vaginal microbiomes are dominated by *Lactobacillus* species, including *L. crispatus*, *L. gasseri, L. jensenii,* and *L. iners* (*13*). *Lactobacillus* spp. produce antimicrobial compounds, such as lactic acid (*14*), hydrogen peroxide (*15*), bacteriocins (*16*), and an arginine deaminase enzyme (*17*), which all may help inhibit the growth of pathogenic bacteria. During BV, this protective microbiome shifts towards a higher abundance of facultative or obligate anaerobic microbes including *Gardnerella* spp., *Prevotella* spp.*, Atopobium* spp.*, Mobiluncus* spp.*, Sneathia* spp., and BV-associated bacteria (BVAB) 1-3 (*18–20*).

*G. vaginalis* is the most well-studied BV-associated anaerobe. Due to its ability to adhere to the vaginal epithelium and tolerate small amounts of oxygen, it is proposed to be an initial anaerobic colonizer that can replace resident *Lactobacillus* species (*21–24*). *G. vaginalis* can use glycogen, a carbon source that is abundant at the vaginal epithelium (*25*, *26*) and degrades the protective mucus layer through the production of sialidases (*27*, *28*). *G. vaginalis* also secretes vaginolysin (VLY), a cytotoxin capable of killing epithelial cells (*29*, *30*). However, not all *G. vaginalis* strains are sialidase-positive and *G. vaginalis* is also found in healthy women (*24*, *31–33*). Therefore *G. vaginalis* may require other species for BV initiation. *Prevotella bivia* for example produces ammonia that stimulates the growth of *G. vaginalis* (*34*). Such synergistic relationships between different vaginal anaerobes with different pathogenic properties most likely drive BV development.

The pathogenic potential of *Prevotella* species in BV is understudied in comparison to *G. vaginalis*. Previous research mainly focused on *P. bivia*, the most commonly isolated *Prevotella* species during BV (*20*). However, recent studies demonstrate that *P. timonensis* is also often found in women with BV (*35–38*). The alternative name *Hoylesella timonensis* was recently proposed for *P. timonensis* (*39*). Due to the high similarity of their 16S rRNA genes, many studies could not discriminate between different *Prevotella* spp. (*40*). Vaginal *Prevotella* spp. in general have been associated with increased cytokine levels in the cervicovaginal fluid (*41–43*). Other reports suggest *Prevotella* spp. may participate in biofilm formation and mucus degradation (*27*, *40*). We have shown that *P. timonensis,* but not *P. bivia,* induces a strong pro-inflammatory response through dendritic cell activation (*44*) and increases HIV-1 uptake by Langerhans cells, turning these cells into HIV-1 reservoirs (*45*). *Prevotella* spp. have also been associated with sialidase activity in vaginal secretions of BV patients (*10*). *P. bivia* has sialidase activity that targets the vaginal mucus layer (*27*) and leads to increased adhesion of other BV-associated bacteria, including *A. vaginae* (*46*)*. P. timonensis* also exhibited sialidase activity and altered mucin expression in the human endometrial epithelial cell line HEC1-A (*40*). However, the role that the different *Prevotella* strains play in BV is currently not clear. In this study, we set out to determine the pathogenic properties of *P. timonensis* compared to *P. bivia* and other BV-associated bacteria, focusing on bacterial interactions with human cells and glycans. We conclude that *P. timonensis* has unique virulence traits that might play an important role during initiation and development of BV.

## Results

### *Prevotella timonensis* adheres to vaginal and endocervical cells

Attachment to the vaginal epithelium is thought to be the first step towards replacement of commensal *Lactobacillus* species and colonization by bacterial anaerobes (Fig. 1A). We investigated the extent to which different BV-associated bacteria can attach to vaginal and endocervical cells and included commensal *L. crispatus* as a control. The vaginal cell line VK2/E6E7 and endocervical cell line End1/E6E7 were grown to a fully confluent monolayer followed by incubation with *L. crispatus, G. vaginalis, P. timonensis,* or *P. bivia* at MOI 10 in anaerobic conditions. After 18 hours, the percentage of attached bacteria was determined by colony counting. We observe that commensal *L. crispatus* adhered well to both VK2/E6E7 and End1/E6E7 cell lines, at 66% and 81% of the total bacterial inoculum, respectively, while *P. bivia* was the least adherent bacterium, with 12% attachment to VK2/E6E7 cells and 2% to End1/E6E7 cells. *G. vaginalis* and *P. timonensis* showed comparable intermediate binding, with adhesion percentages varying between 15% and 40% (Fig. 1B).

**Figure 1.**
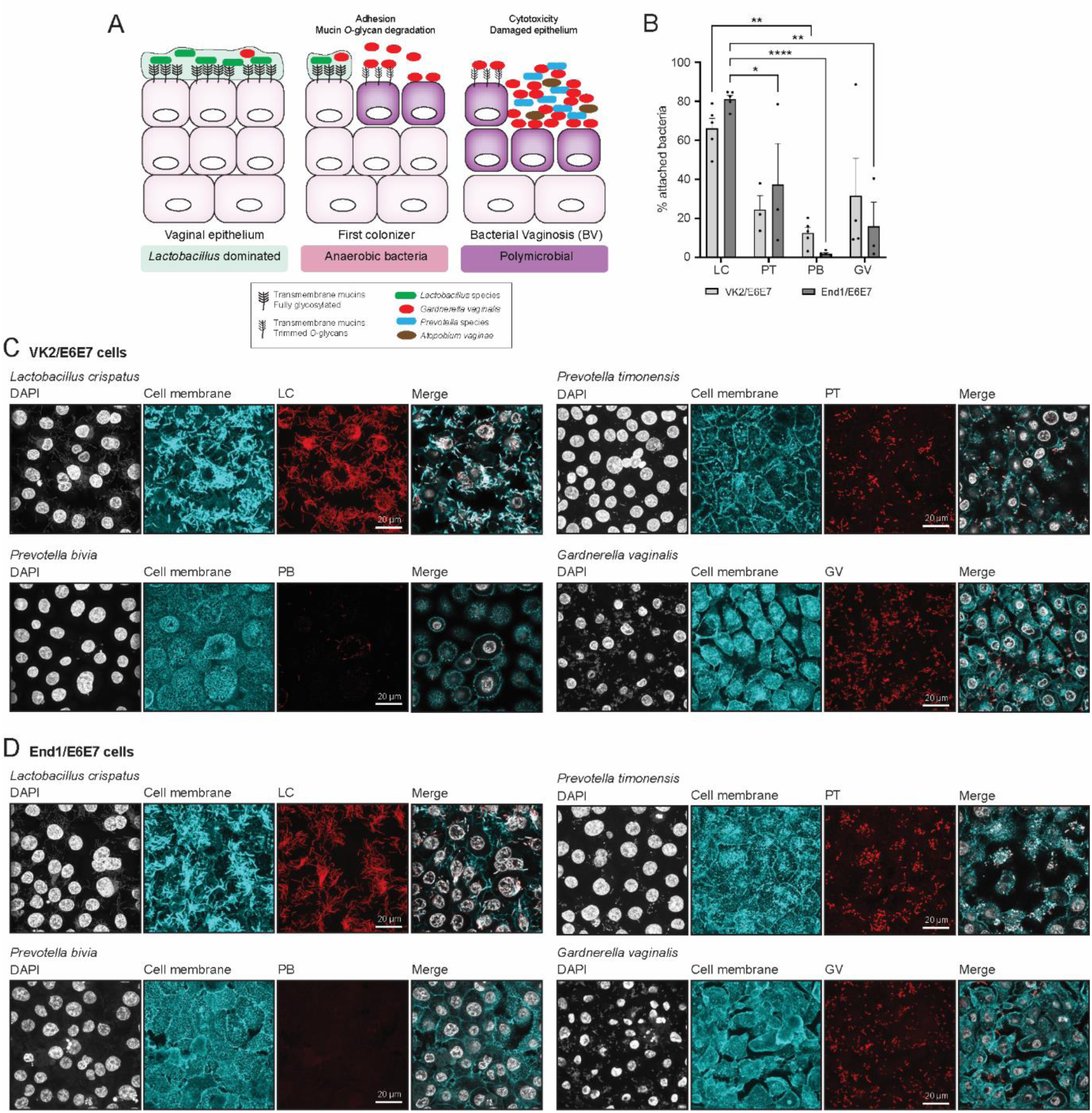
*Prevotella timonensis* can adhere to the vaginal and endocervical epithelium. (**A**) Schematic representation of the different microbial communities of the vaginal epithelium in the healthy state and during the development of bacterial vaginosis. (**B**) Percentage of adhesion of *L. crispatus* (LC), *P. timonensis* (PT)*, P. bivia* (PB), and *G. vaginalis* (GV) to VK2/E6E7 and End1/E6E7 cells assessed by quantification of colony forming units (CFUs). The graph represents the average and SEM of at least 3-4 independent experiments. (**C, D**) FISH in combination with confocal microscopy of *L. crispatus*, *P. timonensis, P. bivia,* and *G. vaginalis* adhesion to (**C**) VK2/E6E7 and (**D**) End1/E6E7 cells stained for WGA and using PNA probes. For each bacterium, the corresponding PNA signal is shown in red, cell surface in cyan (WGA), and DAPI in white. White scale bars represent 20 µM.

In an independent set of experiments, we assessed bacterial binding to cell surfaces by using fluorescence *in situ* hybridization (FISH). We designed specific fluorescently labeled peptide nucleotide acid (PNA) probes for *P. timonensis* (PT-Cy3) and *P. bivia* (PB-Cy3). We used a previously reported PNA probe for *G. vaginalis* (Gard162-AF488) and a general 16S probe for *L. crispatus* (EUB338-AF488). Bacteria were adhered to coated glass slides to test the specificity of the PNA probes and all four probes showed good correlation with the DAPI signal (Fig. S1). We then infected confluent VK2/E6E7 and End1/E6E7 cells with bacteria at a MOI 50 for 18 h anaerobically. The infected epithelial monolayers were stained with the FISH probes and Wheat Germ Agglutinin (WGA) to visualize the epithelial surface. We again found that *G. vaginalis* and *P. timonensis* attached more effectively to the epithelial surface compared to *P. bivia* (Fig. 1C, D). *L. crispatus* showed strong adherence to both cell lines (Fig. 1D, E). The gram-positive *L. crispatus* required a specific permeabilization buffer to achieve efficient labeling of the bacteria with the PNA probe, which led to increased WGA staining of the bacteria in addition to the epithelial cells (Fig. 1C, D). Together, these colony counting and FISH experiments demonstrate that *P. timonensis* can adhere to the surface of vaginal and endocervical monolayers, to comparable levels as the well-known BV-associated pathogen *G. vaginalis*.

### *P. timonensis* does not cause cell cytotoxicity and does not induce major inflammatory responses

We next investigated the cytotoxic and inflammatory potential of *P. timonensis* after adhesion to VK2/E6E7 and End1/E6E7 cell lines. We infected confluent epithelial monolayers with the selected bacteria at a MOI of 10 and 100 for 18 h anaerobically and measured LDH release, an indicator of cellular cytotoxicity. As previously described, *G. vaginalis* was highly cytotoxic, resulting in an LDH release of approximately 70% of the maximum release of the total monolayer (Fig. 2A-B). Incubation with *P. timonensis*, *P. bivia,* or *L. crispatus* for 18 hours did not result in increased LDH release compared to uninfected cells (Fig. 2A-B).

**Figure. 2.**
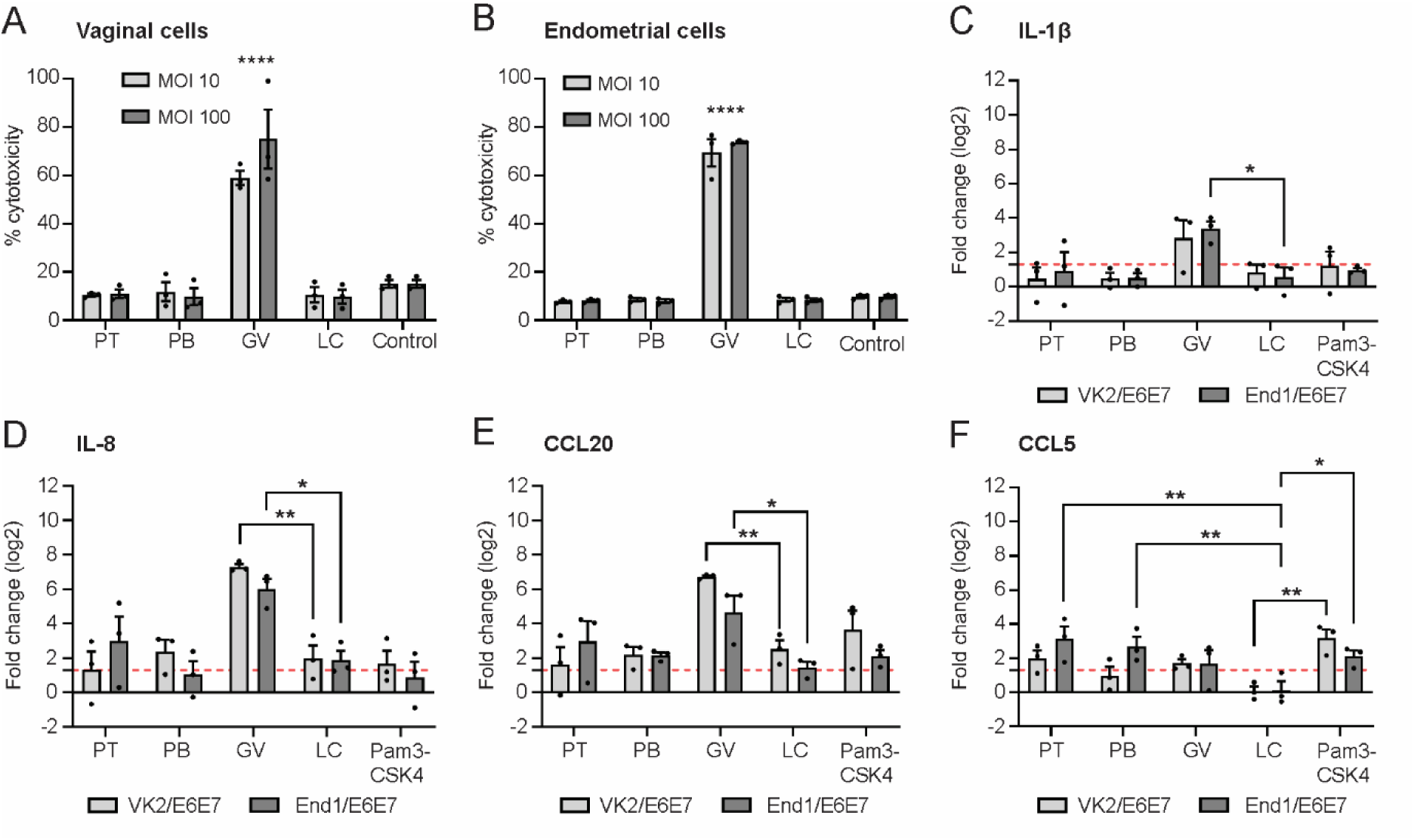
*Prevotella timonensis* does not induce cellular cytotoxicity nor is it highly inflammatory. LDH release of (**A**) VK2/E6E7 and (**B**) End1/E6E7 cells after 18 h infection with *P. timonensis* (PT), *P. bivia* (PB), *G. vaginalis* (GV), or *L. crispatus* (LC) at MOI 10 and 100. (**C-F**) RT-qPCR analysis of VK2/E6E7 and End1/E6E7 cell lines incubated with *P. timonensis* (PT), *P. bivia* (PB), *G. vaginalis* (GV), or *L. crispatus* (LC) at MOI 10 demonstrating expression of (**C**) IL-1β, (**D**) IL-8, € CCL20, and (**F**) CCL5. TMEM222 was used as the reference gene. As a positive control, cells were stimulated with the TLR ligand Pam3CSK4 to induce expression of pro-inflammatory cytokines. The red dotted line marks significant upregulation compared to non-infected cells. The graph represents the average and +/-SEM of at least three independent experiments.

To determine whether the different vaginal bacteria trigger an inflammatory response, we incubated confluent epithelial monolayers with bacteria at MOI 10 and 100 for 18 h anaerobically and measured the mRNA expression of the cytokines IL-1β, IL-8, CCL5, and CCL20 using quantitative RT-PCR.

Compared to the commensal *L. crispatus*, only *G. vaginalis* significantly increased IL-1β, IL-8, and CCL20 expression in both VK2/E6E7 and End1/E6E7 cells (Fig. 2C-E). CCL5/RANTES, a chemoattractant of T lymphocytes and monocytes (*47*) was slightly but significantly upregulated in endocervical cells incubated with *P. timonensis* and *P. bivia*, but not *G. vaginalis* (Fig. 2F). These results suggest that despite its extensive attachment to the epithelial surface, *P. timonensis* does not induce a strong inflammatory response in vaginal or endocervical cells.

### Utilization of glycogen and mucins as carbon sources by BV-associated bacteria

Vaginal and cervical epithelial cells produce high amounts of glycogen, which is deposited onto the epithelium once epithelial cells are shed and lysed (*48*, *49*) and can serve as a carbon source for the resident vaginal bacteria (*50*). We investigated whether our selected BV-associated bacteria could utilize glycogen for growth. Carbohydrates were removed from each bacterium-specific medium and supplemented with 0.5% glycogen. *P. timonensis, P. bivia,* and *G. vaginalis* were all able to grow on glycogen (Fig. 3A-C). Interestingly, *G. vaginalis* reached a higher OD in the basal media supplemented with glycogen compared to the complete specific media, demonstrating a preference for glycogen as carbon source (Fig. 3C). *Akkermansia muciniphila,* a member of the intestinal microbiota known to degrade mucins, did not grow on glycogen (Fig. 3D), supporting the notion that glycogen is a preferred carbon source for vaginal-associated bacteria.

**Figure 3.**
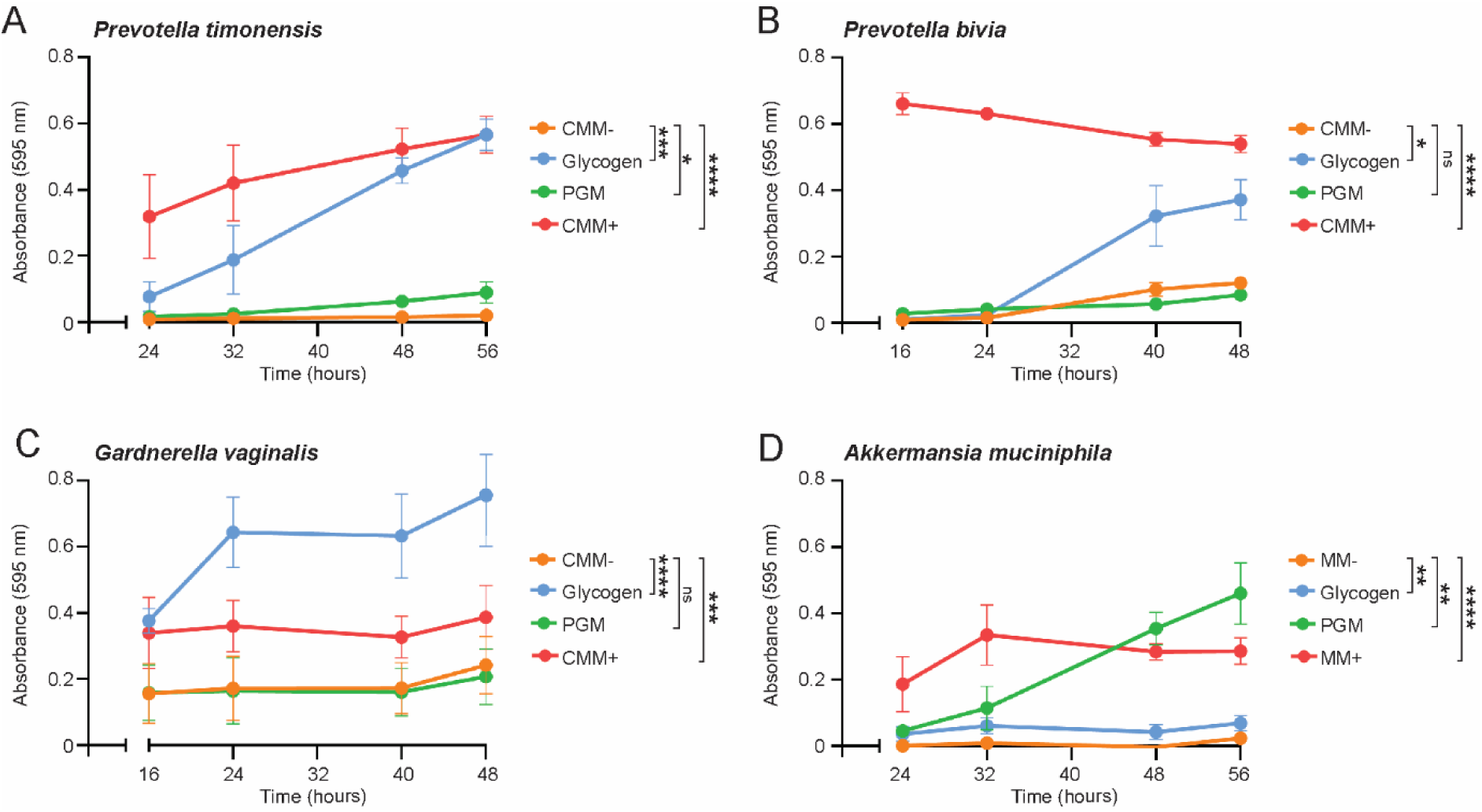
Utilization of glycogen and mucins as carbon sources by BV-associated bacteria. Growth of (**A**) *P. timonensis*, (**B**) *P. bivia*, (**C**) *G. vaginalis,* and mucin-degrader *Akkermansia muciniphila* (**D**), on basal medium without carbohydrates (CMM-or MM-), basal medium supplemented with 0.5% glycogen, basal medium supplemented with 0.5% purified porcine gastric mucins (PGM), or complete medium with carbohydrate (CMM+ or MM+) for up to 56 hours. The graph represents the average and +/-SEM of at least three independent experiments.

The cervicovaginal mucus that covers the vaginal and endocervical epithelium facilitates uterine lubrication and microbial clearance (*51*). We assessed whether the BV-associated bacteria could use mucins as a carbon source by supplementing the basal media with 0.5% purified porcine gastric mucins (PGM). *A. muciniphila* grew well on mucins (Fig. 3D), but *P. bivia* and *G. vaginalis* did not exhibit increased growth on mucins compared to the basal medium without carbohydrates (Fig. 3B, C). *P. timonensis* showed a small but significant increase in growth in the mucin-containing medium compared to the basal medium without carbohydrates (Fig. 3A), suggesting that *P. timonensis* might degrade and utilize mucins.

### The genome of *P. timonensis* predicts a high *O*-glycan degradation potential

To determine the genetic potential of *P. timonensis* and the other BV-associated bacteria to degrade different carbon sources, we sequenced the *P. timonensis, P. bivia, G. vaginalis,* and *A. muciniphila* strains used in this study (sequences deposited in PRJEB67799). The genomes were analyzed for the presence of carbohydrate-active enzymes (CAZymes) using the dbCAN2 meta server pipeline for automated CAZyme annotation (*52*). Only CAZyme genes that were predicted by at least two out of three annotation tools were selected. Detected CAZyme genes included glycoside hydrolases (GH), carbohydrate esterases (CE), glycosyl transferases (GT), carbohydrate-binding modules (CBM), and auxiliary activities (AA). The mucin degrader *A. muciniphila* presented the highest amount of putative CAZy domains (155 ORFs), followed by *P. timonensis* with 104 ORFs, *P. bivia* with 71 ORFs, and *G. vaginalis* with 40 ORFs (Fig. 4A).

**Figure 4.**
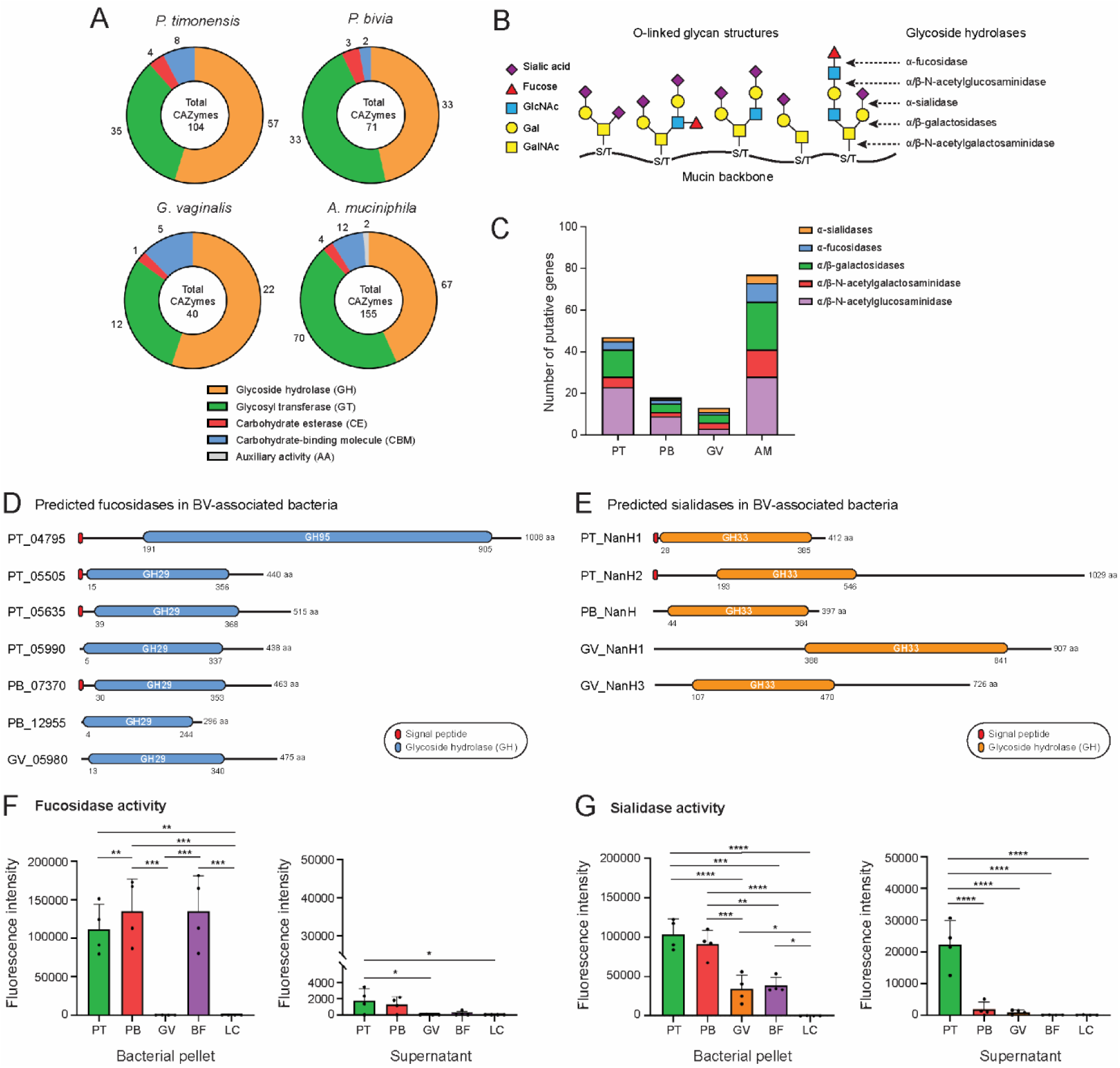
High mucin degradation potential in *Prevotella timonensis*. (**A**) Abundance of predicted carbohydrate-active enzymes (CAZymes) families in the sequenced genomes of our *P. timonensis*, *P. bivia*, *G. vaginalis* and *A. muciniphila* strains. (**B**) Schematic representation of a mucin glycoprotein molecule with protein backbone and diverse *O*-glycan structures. Target sites for different classes of glycosyl hydrolases are depicted. (**C**) Number of identified *O*-glycan-targeting CAZymes in the genomes of the sequenced *P. timonensis* (PT), *P. bivia* (PB), *G. vaginalis* (GV), and *A. muciniphila* (AM) strains. (**D-E**) Domain architecture of the predicted (**D**) fucosidases and (**E**) sialidases of the designated bacteria. The displayed domains are identified by HMMER, Diamond, and Signal IP 6.0 tools and drawn to scale. (**F**) Fucosidase and (**G**) sialidase activities measured in bacterial pellets and supernatants of the different BV-associated bacteria and *B. fragilis* as positive control. Abbreviations: PT (*P. timonensis*), PB (*P. bivia*), GV (*G. vaginalis*), BF (*B. fragilis*), and LC (*L. crispatus*). The graph represents the average and +/-SEM of four independent experiments.

We next examined these results to identify candidate enzymes for degradation of specific substrates. Mucins have polypeptide backbones that are decorated by complex *O*-linked glycan structures that require sequential degradation by glycoside hydrolases with high specificity (Fig. 4B). Within the glycoside hydrolase category, several genes encoding predicted sialidases (GH33 class) and fucosidases (GH29 and GH95 classes) were detected in the genomes of all four bacteria (Fig. 4C). Furthermore, *P. timonensis* and *A. muciniphila* possessed a great number of predicted α/β-galactosidases, α/β-N-acetylgalactosaminidases, and α/β-N-acetylglucosaminidases, enzymes that hydrolyze the glycosidic linkages underlying the terminal sialic acids and fucoses (Fig. 4C). Many of these putative CAZymes contained a signal peptide, suggesting that the proteins may be translocated to the bacterial surface or secreted into the environment (Fig. 4C).

Sialic acids and fucoses cap mucin *O*-glycan structures and are the first monosaccharides that need to be removed for further mucin degradation (*53*). The fucosidase family consists of the retaining fucosidases (GH29) and inverting fucosidases (GH95). The *P. timonensis* genome uniquely encoded a predicted GH95 enzyme in addition to three GH29-containing fucosidases. The *P*. *bivia* and *G. vaginalis* genomes encoded two and one predicted GH29 enzymes, respectively (Fig. 4D). The *P. timonensis* genome encoded two predicted sialidases with a GH33 domain and signal peptides with different domain structures. The NanH1 sialidase is predicted to be 412 amino acids in length and NanH2 is a much larger protein with 1029 amino acids. Both *P. timonensis* sialidases are likely secreted enzymes as they have predicted signal peptides. In an accompanying paper, Pelayo et al., biochemically characterize these two *P. timonensis* sialidases, establishing their activity. The *G. vaginalis* genome encoded two predicted sialidases (NanH1 and NanH3, NanH2 was not present in our *Gardnerella* strain) and a single GH33 sialidase (NanH) was predicted for *P. bivia* (Fig. 4E). Earlier studies showed that *G. vaginalis* NanH2 and NanH3, but not NanH1, had high activity towards 4-methylumbelliferyl *N*-acetyl-α-D-neuraminic acid (4-MU-Neu5Ac) and bovine submaxillary mucin (*54*). These studies underpin the importance of studying enzyme activity. Altogether, our genomic analysis suggests that *P. timonensis* has a larger repertoire of potential mucin-degrading enzymes compared to the other BV-associated bacteria including multiple fucosidases and sialidases.

### *P. timonensis* displays high fucosidase and sialidase activity on the bacterial surface and in the supernatant

Sialidase and fucosidase activity in bacteria are often associated with pathogenic behavior as the removal of terminal monosaccharides from the mucin *O*-glycan structure promotes further degradation by exposing underlying glycans and the mucin peptide backbone that is sensitive to proteases (*53*). To assess the presence of sialidase and fucosidase activities in our vaginal bacterial strains we performed culture-based assays. We also included *Bacteroides fragilis* as a positive control, an intestinal bacterium that is sometimes associated with vaginitis (*55*) and pelvic inflammatory disease (*56*, *57*), and is known to have sialidase and fucosidase activity. Bacteria were grown overnight followed by centrifugation to separate the pellet from the supernatant fraction. To determine fucosidase and sialidase activities, both fractions were incubated with fluorescent substrates and the produced fluorescence by each enzyme was measured.

No fucosidase activity was detected for *G. vaginalis* and *L. crispatus*, which was surprising as the *G. vaginalis* genome does encode a GH29 fucosidase (Fig. 4D). *P. timonensis, P. bivia,* and *B. fragilis*, all displayed high fucosidase activity in the bacterial pellet. In addition, fucosidase activity was detectable in the supernatants of *P. timonensis* and *P. bivia*, but only reached statistical significance in the case of *P. timonensis* compared to the fucosidase-negative *G. vaginalis* supernatant (Fig. 4F). For sialidase activity, the highest cell-bound activity could be measured for *P. timonensis* and *P. bivia* followed by *G. vaginalis* and *B. fragilis*. Both *P. timonensis* and *P. bivia* sialidase activities were significantly higher than those of *G. vaginalis.* Of the supernatant fractions, only that of *P. timonensis* contained detectable sialidase activity, suggesting that this bacterium secreted sialidases into the medium under the conditions tested (Fig. 4G).

### *P. timonensis* sialidase and fucosidase activity leads to *O*-glycan degradation at the vaginal surface

Next, we determined the *O*-glycan-degrading capacity of vaginal bacteria at the vaginal epithelial surface. Vaginal VK2/E6E7 monolayers were incubated with bacteria for 18 h and stained with lectins to detect different mucin glycan structures including fucoses (UEA-1), α2,3 sialic acids (MAL-II), and α2,6 sialic acids (SNA). Visualization by confocal microscopy demonstrated that all glycan structures were present on the vaginal epithelial surfaces in the absence of bacteria (Fig. 5A-D, top panels). UEA-1 staining was significantly reduced after incubation with *P. timonensis* and *G. vaginalis* demonstrating removal of fucose residues, but not after incubation with *P. bivia* and *L. crispatus* (Fig. 5A, D). This result indicated that the *G. vaginalis* fucosidase is active and is perhaps induced after interaction with vaginal epithelial cells. Incubation with *P. timonensis* significantly decreased the staining for α2,3 sialic acids and α2,6 sialic acids on the vaginal epithelial surface (Fig. 5B-D). Reduction of MAL-II, but not SNA, staining was also observed after incubation with *G. vaginalis*, but to a lesser extent than *P. timonensis* (Fig. 5B-D). *P. bivia* and *L. crispatus* did not significantly reduce sialic acid staining (Fig. 5B-D). Overall, these results show that *P. timonensis* efficiently removes sialic acids and fucoses from the vaginal epithelium.

**Figure 5.**
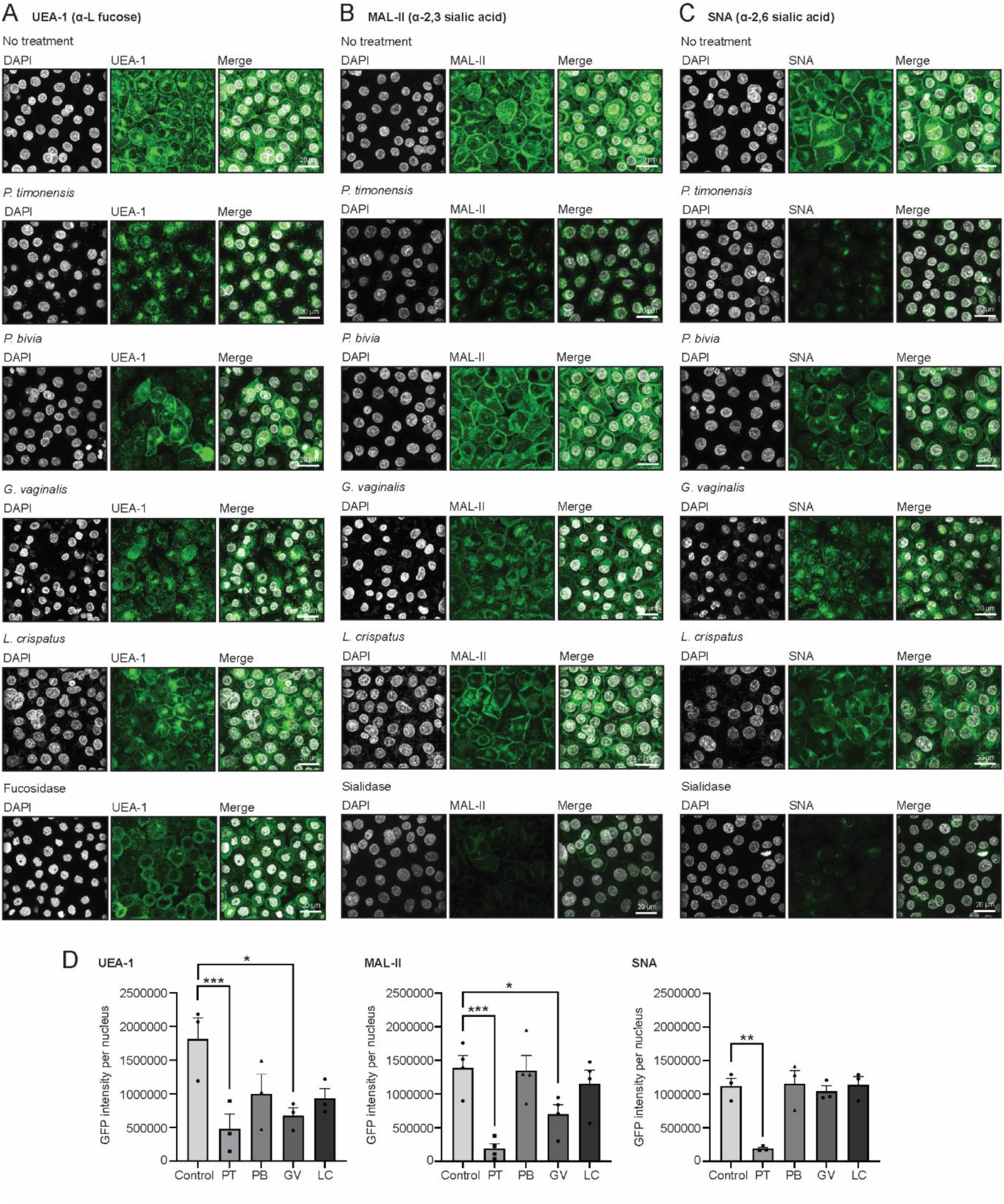
*Prevotella timonensis* effectively removes glycans from the vaginal epithelial surface. Fluorescence confocal microscopy images of mucin *O*-glycan structures after incubation with *P. timonensis*, *P. bivia*, *G. vaginalis*, or *L. crispatus* at a MOI 10 for 18 hours anaerobically. Neuraminidase A and L-fucosidase were added for 3 h as positive controls for sialidase and fucosidases activity. (**A**) UEA-1 (α-L fucoses), (**B**) MAL-II (α-2,3 sialic acids), and (**C**) SNA (α-2,6 sialic acids) stainings are shown in green, and DAPI in white. White scale bars represent 20 µM. (**D**) Quantification of UEA-1, MAL-II, and SNA stainings from figure 5A-C. The graph represents the average and +/-SEM of at least three independent experiments.

To investigate if the two *P. timonensis* sialidases (PtNanH1 and PtNanH2) could remove sialic acids from the epithelial surface, we incubated the vaginal monolayers with recombinantly expressed and purified PtNanH1 and PtNanH2. Incubation with either enzyme led to a significant reduction of both MAL-II and SNA (Fig. 6A-C). Next, vaginal epithelial monolayers were incubated with *P. timonensis* in the presence of either the broad sialidase inhibitor DANA or Zanamivir, an inhibitor that was found in the accompanying study to be effective toward *P. timonensis* sialidases. In the presence of DANA or Zanamivir, removal of α2,3 sialic acids and α2,6 sialic acids from the vaginal epithelial surface by *P. timonensis* was significantly reduced, demonstrating efficient inhibition of the bacterial sialidases and highlights the role of these enzymes in glycan degradation (Fig. 6D-F). In conclusion, the BV-associated bacterium *P. timonensis* has a high potential for *O*-glycan degradation at the vaginal epithelial mucosal surface through a diverse array of glycogen hydrolases including highly active sialidases.

**Figure 6.**
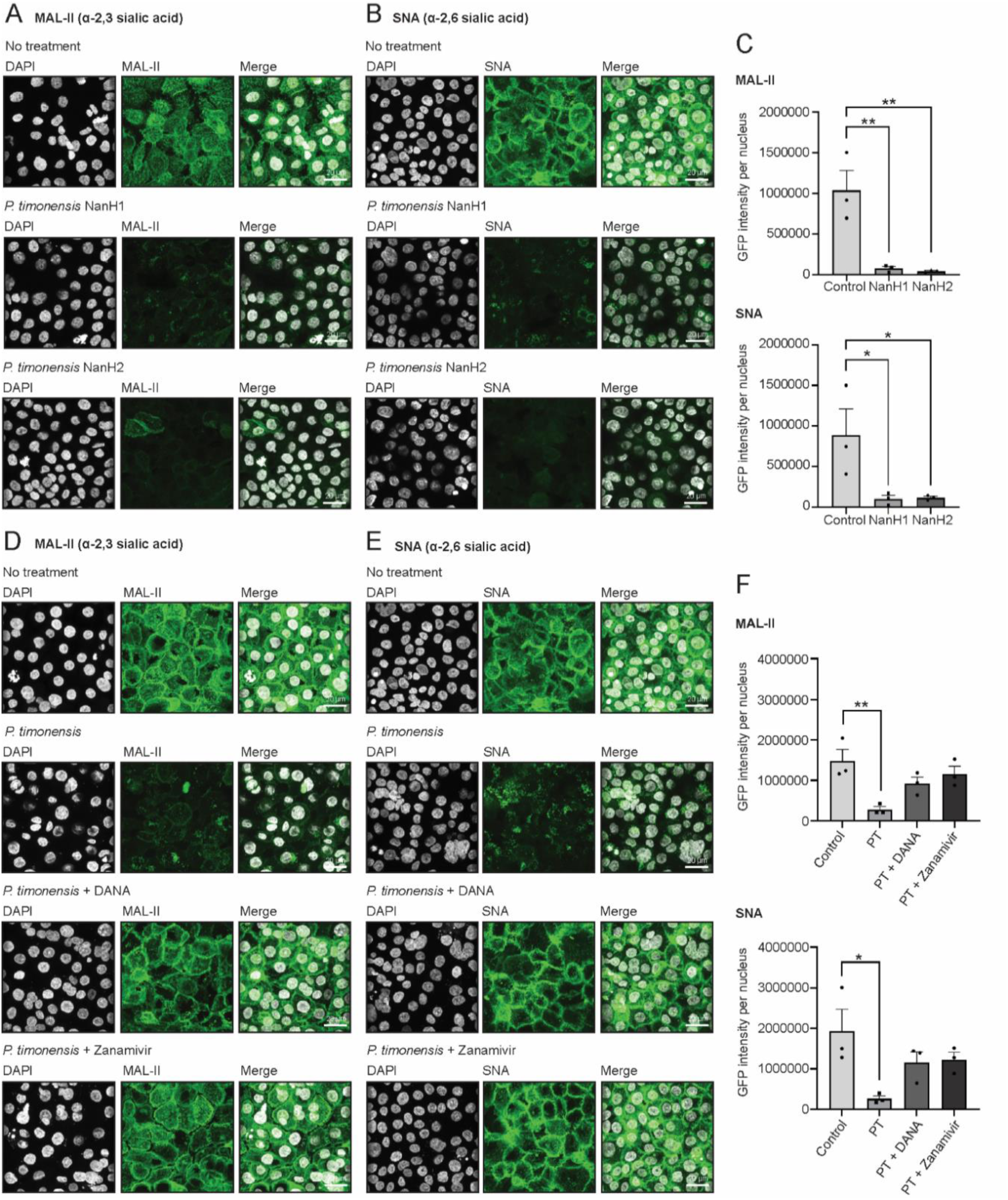
*P. timonensis* sialidase activity at the vaginal mucosal surface can be inhibited by the sialidase inhibitors DANA and Zanamivir. Fluorescence confocal microscopy images of sialic acid staining after incubation with 1 µM of recombinant *P. timonensis* sialidases NanH1 and NanH2 for 3 hours anaerobically. (**A**) MAL-II (α-2,3 sialic acids) and (**B**) SNA (α-2,6 sialic acids) stainings are shown in green and DAPI in white. White scale bars represent 20 µM. (**C**) Quantification of MAL-II and SNA stainings from figure 6A-B. Fluorescence microscopy images of (**D**) MAL-II (α-2,3 sialic acids) and (**E**) SNA (α-2,6 sialic acids) after *P. timonensis* infection at a MOI 10 for 18 hours anaerobically in the presence/absence of 1 mM DANA or Zanamivir. Lectin stainings are shown in green and DAPI in white. White scale bars represent 20 µM. (**F**) Quantification of MAL-II and SNA stainings from figure 6D-E. The graph represents the average and +/-SEM of three independent experiments.

## Discussion

BV is one of the most common pathological conditions in women of different ages that increases susceptibility to sexual transmitted infections and negatively impacts fertility and quality of life. Unlike the health-associated vaginal microbiome, which is dominated by *Lactobacillus* species, BV is characterized by a polymicrobial infection of different anaerobes including *G. vaginalis, A. vaginae* and different *Prevotella* species (Fig. 1A). High sialidase activity can be detected in the vaginal discharge of women with BV (*10–12*) and persistence of sialidase-positive bacteria is a risk factor for subclinical intrauterine infections and preterm birth (*58*).

Thus far, *G. vaginalis* and *P. bivia* were considered to be the main producers of sialidases in the cervicovaginal environment (*59–62*). In this study and the accompanying paper by Pelayo et al., we demonstrate that *P. timonensis* has high sialidase activity and should be considered amongst the bacteria that play a pivotal role in the initiation and progression of BV. We found that *P. timonensis* had the highest sialidase activity of the BV-associated bacterial strains tested. In addition to cell-bound sialidase activity, *P. timonensis* was the only bacterium with detectable secreted sialidase activity (Fig. 4G). After attachment to vaginal epithelial cells, *P. timonensis* removed the majority of surface α2-3-linked and α2-6-linked sialic acids (Fig. 5B-C), and the two identified *P. timonensis* sialidases (PtNanH1 and PtNanH2) were highly active at removing sialic acids from the vaginal epithelial cell surface (Fig. 6 D-F). Notably, the sialidase activity of *P. timonensis* at the vaginal epithelial surface could be blocked with DANA and Zanamivir inhibitors (Fig. 6A-C). In addition to sialidase activity, *P. timonensis* also displayed fucosidase activity in culture and during attachment to the vaginal epithelium (Fig. 4F, 5A). Sialidases and fucosidases are essential enzymes that can initiate degradation of mucin O-glycan structures of secreted and epithelium-bound mucins. The removal of sialic acids renders mucins more vulnerable to further degradation by glycosyl hydrolases and proteases (*63*). A recent paper showed that recombinant sialidases of *Gardnerella* species led to desialylation of glycans in VK2/E6E7 and induced pathways of cell death, differentiation, and inflammatory responses (*64*). Therefore, these enzymes are important virulence factors that can contribute to the establishment and development of BV.

Investigating bacterial nutritional preference for cervicovaginal mucus and glycogen is important to understand how different members of the vaginal microbiome thrive in this unique environment. Sialidases and fucosidases are crucial for bacterial growth on mucin (*65*). Besides these enzymes, *P. timonensis* also encoded a wide array of other mucin-degrading enzymes (Fig. 4A, C) and showed a small but significant growth on mucins as the sole carbon source (Fig. 3A-D). *G. vaginalis* and *P. bivia* encode fewer mucin-degrading enzymes (Fig. 4A, C) and did not grow in mucin-enriched media (Fig. 3B-C). For these experiments, we used pig gastric mucus (PGM) containing 5-N-glycolylneuraminic acid (Neu5Gc) (*66*). This mucus might be less suitable for human microbiota as human mucus does not contain Neu5Gc and it was previously suggested that *G. vaginalis* is not capable of degrading Neu5Gc (*61*). Therefore, future experiments should be conducted with human (vaginal) mucus to conclusively establish the growth capacities of the different vaginal microbiota on cervicovaginal mucus. Glycogen is a large, highly branched D-glucose polymer that is abundant in vaginal tissue (*48*, *67*) but present at reduced levels in women with BV (*68–70*). Several *Lactobacillus* spp. have been shown to directly use glycogen for growth (*71*, *72*). Here we show that *P. timonensis*, *G. vaginalis,* and *P. bivia* were all able to grow on glycogen as single nutrient source, which is in line with the presence of α-glucosidases in their genomes (*25*, *50*). Overall, glycogen utilization seems to be a shared trait of vagina-associated bacteria, indicating their adaptation to the vaginal environment.

Adhesion to the cell epithelium is a crucial step in BV and many studies in the field indicate a stepwise disease progression with primary and secondary bacterial colonizers (Fig. 1A). *G. vaginalis* is an important primary colonizer as this bacterium can adhere to the vaginal epithelium and potentially form a biofilm (*21–24*). Other anaerobic bacteria such as *P. bivia* and *A. vaginae,* can join the *G. vaginalis* biofilm as secondary colonizers (*46*, *73*, *74*). In the current study, we demonstrate that *P. timonensis*, similar to *G. vaginalis* but unlike *P. bivia*, can efficiently bind to both vaginal and endocervical cells (Fig. 1B, D, E). Previously, it has been shown that *P. timonensis* can induce elongated microvilli in a 3D endometrial epithelial cell model, and it was speculated that these changes might induce increased adhesion of this species and of other secondary colonizers (*40*). Based on these combined observations, we propose that *P. timonensis* may be an initial colonizer of the vaginal epithelium and does not require an established *G. vaginalis* biofilm. After attachment, the high sialidase and fucosidase activity of *P. timonensis* at the vaginal epithelial surface removes the protective terminal glycans of the glycocalyx likely creating new binding sites for secondary colonizers (*64*, *75*) and enhancing bacterial colonization of the upper parts of the FRT (*76*).

While *P. timonensis* is perhaps an initial colonizer during BV, it does not contribute to cytotoxicity nor did it induce a pro-inflammatory response in a similar manner to *G. vaginalis*. Only *G. vaginalis* induced high LDH release by both vaginal and endocervical cells while *P. timonensis, P. bivia,* and *L. crispatus* were not cytotoxic (Fig. 2A, B). To induce cytotoxicity, *G. vaginalis* expresses the cytotoxin vaginolysin (vly) (*29*, *30*) and also utilizes membrane vesicles (*77*). Cytotoxicity might be an important aspect of *G. vaginalis* virulence, as strains isolated from women with BV were more cytotoxic than non-BV isolates (*33*). In our infection experiments with single species of bacteria, *G. vaginalis* strongly induced expression of IL-1β, IL-8, or CCL20 in vaginal and endocervical epithelium while *P. timonensis*, *P. bivia,* and *L. crispatus* did not significantly induce pro-inflammatory cytokines (Fig. 2C-E), which was in line with previous reports that investigated single species (*40*, *78*, *79*). Therefore, the current data suggest that *P. timonensis* by itself is not a promoter of BV-associated inflammation as has been observed for *G. vaginalis* and *A. vaginae* (*79*, *80*). Because *in vivo* observations in cervicovaginal fluid indicate increased cytokine levels when *Prevotella* spp. are present in the vagina (*41*, *42*), the contributions of different *Prevotella* species to pro-inflammatory responses in more complex polymicrobial infections remain to be established.

This study provides evidence that the understudied vaginal bacteria *P. timonensis* has pathogenic properties that could support primary colonization of the female reproductive tract in BV. Unlike *G. vaginalis,* the virulence traits of *P. timonensis* do not include cell cytotoxicity nor triggering of a strong pro-inflammatory response but rather a strong and previously unappreciated capacity to degrade the protective epithelial mucus layer through sialidase and fucosidase activity. We also demonstrate that the *P. timonensis* sialidase activity at the vaginal epithelial glycocalyx can be efficiently inhibited by small molecule inhibitors. For *G. vaginalis,* it was previously demonstrated that a sialidase inhibitor also reduced cellular invasion (*81*). The application of sialidase inhibitors in BV treatment might therefore be an interesting novel therapeutic approach to reduce bacterial adhesion, invasion and mucosal damage.

## Materials and Methods

### Cell lines, bacterial strains, and culture conditions

VK2/E6E7 (ATCC, CRL-2616) and End1/E6E7 (ATCC, CRL-2615) cells were routinely grown and maintained as indicated in the supplementary materials. The bacterial strains used in this study are listed in Table 1. Bacterial media and growing conditions can be found in the supplementary materials.

**Table 1.** Overview of bacterial strains used in this study.

| Species | Strain | Isolation site | Growth medium | Plate medium | Taxonomy ID | Reference |
| --- | --- | --- | --- | --- | --- | --- |
| *Prevotella timonensis* (*Hoylesella timonensis*) | CRIS 5C-B1 | Human vagina | CMM | Chocolate, bioTRADING (K516P090KP) | 679189 | BEI resources, n. d. |
| *Prevotella bivia* | DSM 20514 | Endometrium | CMM | NYC | 868129 | DSMZ |
| *Gardnerella vaginalis* | CCUG 72422 | Human vagina | NYC | NYC | n/a | CCUG |
| *Lactobacillus crispatus* | RL10 | Human vagina | MRS | MRS | n/a | NCCB 100715 |
| *Bacteroides fragilis* | ATCC 25285 | Appendix abscess | NYC | NYC | n/a | ATCC |
| *Akkermansia muciniphila* | IMS 1-22 | Human feces | MM | MM | n/a | In house |

### Adhesion assay

VK2/E6E7 and End1/E6E7 cells were seeded in a 12-well plate and grew until full confluency. Cells were infected with overnight bacterial cultures at a MOI of 10 for 18 h in anaerobic conditions. Serial dilutions from the supernatant and cell suspensions were plated in their specific plate media. Colonies were counted to calculate the percentage of adherent bacteria, as described in the supplementary methods.

### Peptide nucleotide acid (PNA) probe in-silico design

PNA probes for the specific detection of *P. timonensis* or *P. bivia* were designed using the protocol described in detail in the supplementary materials. The resulting PNA probes were named PT-Cy3 and PB-Cy3. The Gard162-AF488 (*82*) and EUB338-AF488 (*83*) PNA probes used in this study have been previously described. All probes are listed in Table 2.

**Table 2.**
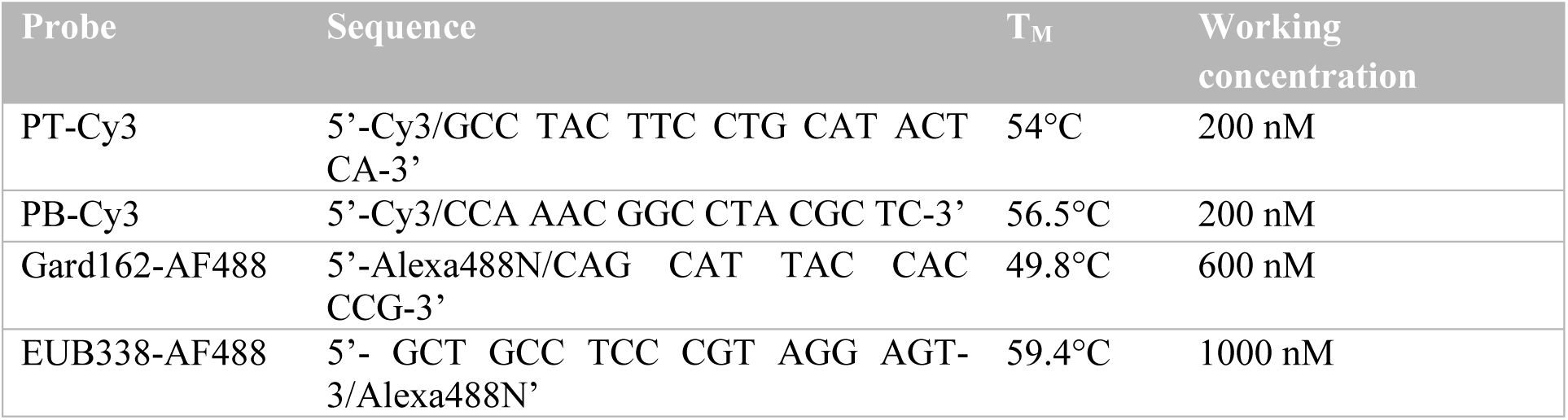
Sequences of the PNA/FISH probes.

### Fluorescence In Situ Hybridization (FISH) and confocal microscopy

Briefly, confluent monolayers of VK2/E6E7 and End1/E6E7 cells were infected with *G. vaginalis, P. timonensis,* or *P. bivia* at a MOI of 50 for 18 h at 37°C in anaerobic conditions. Cells were washed and stained with Wheat Germ Agglutinin-663 (WGA-633, Invitrogen, W21404). Then, cells were fixed and stained with 1000 nM EUB338-AF488 probe, 600 nM Gard162-AF488 probe, 200 nM PT-Cy3, or 200 nM PB-Cy3 probe in hybridization buffer for 2 h at 50°C in a humidity chamber. Slides were washed, stained with DAPI, and mounted for imaging on a Leica SPE-II confocal microscope. Additional details of the FISH staining protocol can be found in the supplementary materials.

### Lectin staining and confocal microscopy

For infection experiments and *O*-glycan analysis, epithelial cells were grown and infected with bacteria as described under the FISH protocol. Cells were treated with 200 U/mL of α2,3,6,8,9 neuraminidase A (NEB Bioke, P0722L) and 0.6 U of α1,2,3,4,6-L-fucosidase (Megazyme, E-FUCHS) for 3 h as positive controls for sialidase and fucosidase activity. Cells were incubated with lectins Sambucus Nigra Lectin biotinylated (SNA) at 1:200, Maackia Amurensis Lectin II biotinylated (MAL-II) at 1:100, and Ulex Europaeus Agglutinin I (UEA-1) at 1:100, followed by incubation with Streptavidin-488 at 1:100 and DAPI at 1:1000. Slides were washed and mounted for imaging on a Leica SPE-II confocal microscope.

### Cytotoxicity assays

VK2/E6E7 and End1/E6E7 cells were grown until full confluency in 96-well plates. Overnight cultures of *P. timonensis, P. bivia, G. vaginalis,* and *L. crispatus* were used to infect the cells at a MOI of 10 and 100 for 18 h under anaerobic conditions. The presence of released LDH in the supernatant was assessed using the Cytotox 96 Non-Radioactive Cytotoxicity Assay (Promega, G1780). The extended protocol can be found in the supplementary materials.

### Reverse transcription quantitative polymerase chain reaction (RT-qPCR)

Non-confluent VK2/E6E7 and End1/E6E7 cells were infected with *G. vaginalis, P. timonensis,* or *P. bivia* at a MOI of 10 for 18 h at 37°C in anaerobic conditions. RNA was extracted and treated with DNAse I. Primers used in the RT-qPCR reactions can be found in Table 3. All cycle thresholds were averaged from triplicate reactions and normalized to the housekeeping gene TMEM222. Fold changes were calculated using the delta-delta Ct method. Additional details of bacterial infection and RT-qPCR protocols can be found in the supplementary materials.

**Table 3.**
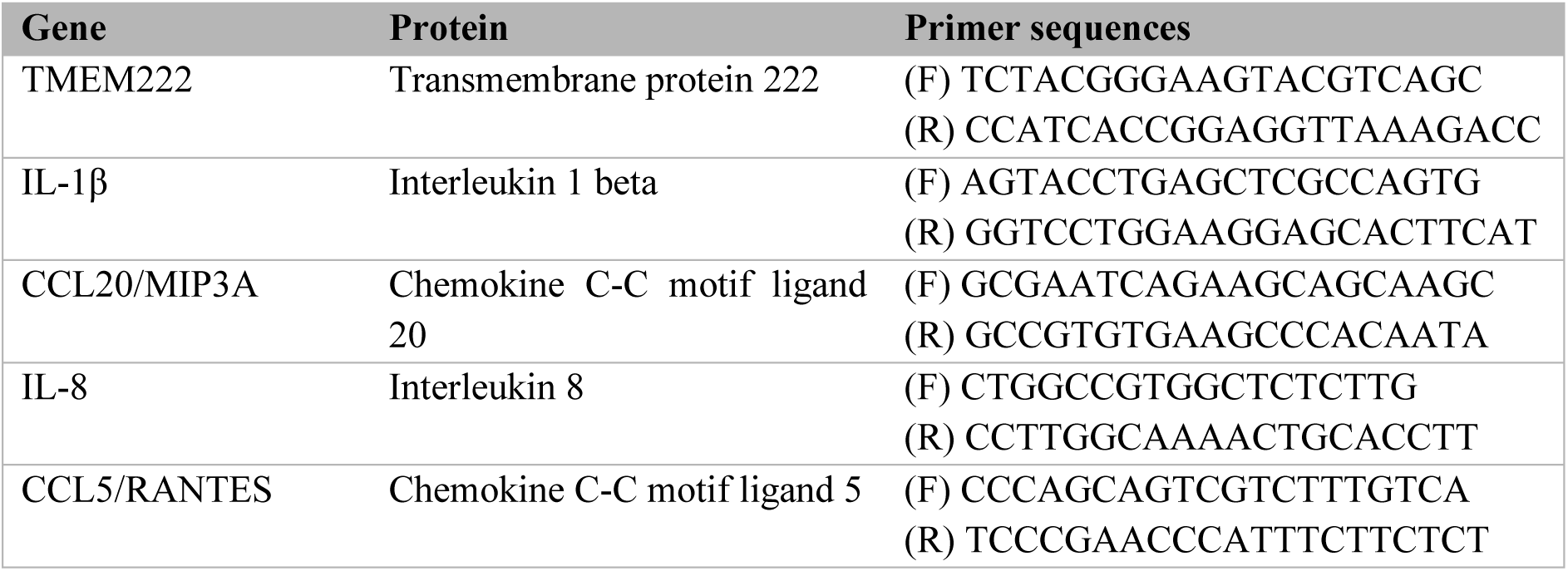
Primer sequences for RT-qPCR. F: forward primer; R: reverse primer.

### Mucin and glycogen growth assays

Mucins were purified from commercially available porcine gastric mucins (PGM, Sigma-Aldrich, M2378) as described in the supplementary materials. Bacterial cultures of *P. timonensis, P. bivia, G. vaginalis,* and *A. muciniphila* were diluted to OD600 = 0.02 in their respective medium without carbohydrates. In a 24-well plate, diluted bacterial cultures were mixed 1:1 with their corresponding medium without carbohydrates, supplemented with purified mucins or glycogen (Sigma-Aldrich, 10901393001) at a final concentration of 0.5% w/v. As a positive control, complete medium with carbohydrates was used. The cultures were incubated in anaerobic conditions for up to 56 h at 37°C. During incubation, absorbance of 100 µL of each culture was measured at 595 nm with the FLUOstar Omega microplate reader at 24, 32, 48, and 56 h for *P. timonensis* and *A. muciniphila*, and at 16, 24, 40, and 48 h for the faster-growing bacteria *P. bivia* and *G. vaginalis*.

### Bacterial whole genome sequencing and CAZyme analysis

Bacterial DNA was isolated and sequenced using Nanopore technology. Detailed information can be found in the supplementary materials. Predicted bacterial protein sequences were used to analyze the presence of carbohydrate-active enzymes using the CAZy database and dbCAN3 meta server. CAZymes identified with at least two out of three tools (HMMER: dbCAN, DIAMOND: CAZy, and HMMER: dbCAN_sub) were considered for further analysis.

### Enzymatic activity assays

Fucosidase and sialidase activities were measured in overnight bacterial pellets and concentrated supernatants using fluorogenic substrates 4-Methylumbelliferyl α-L-fucopyranoside and 4-Methylumbelliferyl N-acetyl-a-D-neuraminic acid sodium salt, respectively. The detailed enzyme assay protocols can be found in the supplementary materials.

### Cloning, heterologous expression, and purification of *P. timonensis* sialidase genes

Full-length sialidase genes were amplified from genomic DNA purifications using primer pairs depicted in Table 4. The resulting gene products were assembled into pET28a expression vector and transformed into DH5α *Escherichia coli* chemically competent cells. Confirmed plasmids were transformed into *E. coli* BL21 (DE3) for protein expression. A more extensive detailed protocol can be found in the supplementary materials.

**Table 4.**
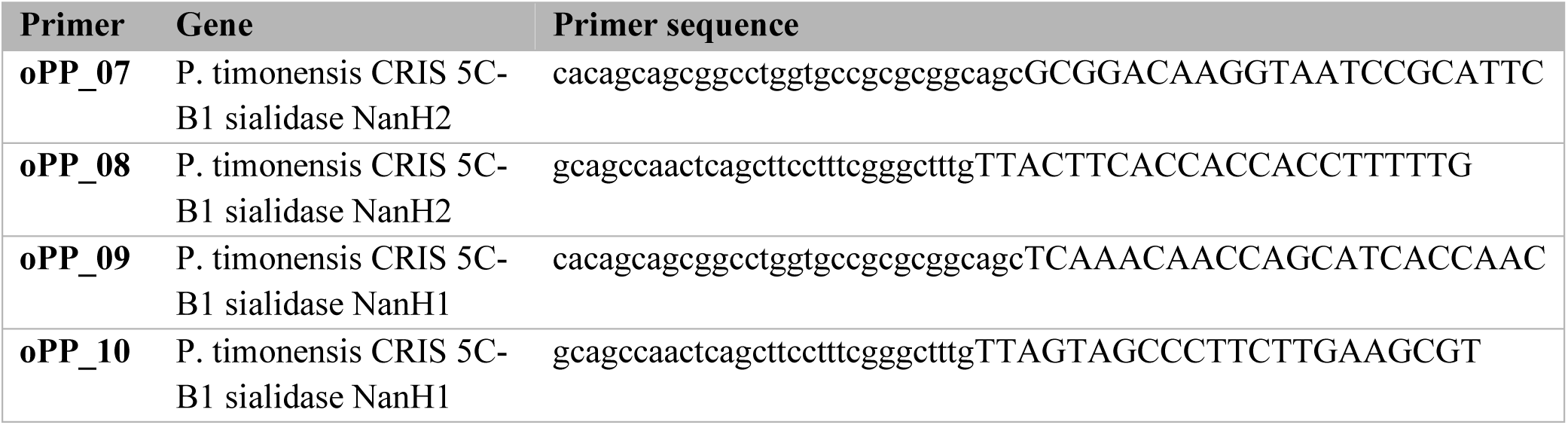
Primer sequences for the cloning of P. timonensis sialidase genes.

## Acknowledgements

We thank Prof. dr. ir. Remco Kort (Vrije Universiteit, the Netherlands) for providing the *L. crispatus* strain and the group of Prof. dr. Piet Cools (University Ghent, Belgium) for providing the *G. vaginalis* strain and for expert advice. We thank Jos P.M. van Putten for advice and support throughout the project. This research is supported by a ZonMW TOP grant that was awarded to K. Strijbis and T. Geijtenbeek (grant number 91218017). K. Strijbis has received funding from the European Research Council (ERC) under the European Union’s Horizon 2020 research and innovation program (ERC-2019-STG 852452).

