## Supplementary materials for "*Prevotella timonensis* degrades the vaginal epithelial glycocalyx through high fucosidase and sialidase activities"

### Cell lines and culturing

VK2/E6E7 (ATCC, CRL-2616) and End1/E6E7 (ATCC, CRL-2615) cells were routinely grown in 75 $cm^2$ flasks at 37°C in 5% $CO_2$ in keratinocyte serum-free (KSF) medium (Gibco, 17005042) supplemented with 0.05 mg/mL bovine pituitary extract, 0.1 ng/mL recombinant human epidermal growth factor, and 0.4 mM $CaCl_2$. Cells were passaged with 1:6 dilution before reaching 80% confluency. For infection assays, cells were seeded at 70% confluency and grown for 4 days until full confluency. The media was refreshed every two days until the day of infection.

### Bacterial strains and culture conditions

The bacterial strains used in this study are listed in Table 1. All bacteria were grown at 37°C under anaerobic conditions (5% $H_2$, 10% $CO_2$, 85% $N_2$) in a Coy Lab's Vinyl Anaerobic Chamber. *P. timonensis* and *P. bivia* were grown in Cooked Meat Medium (CMM, CM0081) supplemented with 1 mg/L vitamin K1 (Sigma, V3501) and 5 mg/L hemin (Sigma, 51280) following the DSMZ #110 protocol (https://www.dsmz.de/microorganisms/medium/pdf/DSMZ_Medium110.pdf). *G. vaginalis*, and *B. fragilis* were grown in New York City (NYC) medium comprising 4 g/L HEPES (ThermoFisher, 15630-080), 15 g/L peptone (ThermoFisher, 211677), 5 g/L NaCl (VWR, 1064041000,), 5 g/L glucose (Merck, 1083370250), 25 g/L yeast extract (Gibco, 212750), and 10% heat-inactivated FCS (Sigma, F7524). *A. muciniphila* was cultured in Mega Medium (MM) (*1*), and *L. crispatus* in De Man, Rogosa, and Sharpe (MRS) medium (Millipore, 69966).

### Adhesion assay

Overnight bacterial cultures were washed with Dulbecco's Phosphate Buffered Saline (DPBS, Sigma, D8537) and resuspended in KSF medium. VK2/E6E7 and End1/E6E7 cells were seeded in a 12-well plate at 70% confluency and grown for 4 days until full confluency. The cells were then transferred to the anaerobic chamber, washed twice with anaerobic DPBS to remove remaining oxygen, and infected with bacteria at a MOI of 10. After 18 h incubation at 37°C, the supernatant was transferred to a new eppendorf tube. Cells were washed 3 times with DPBS, incubated with 500 µL 0.25% trypsin (ThermoFisher, 25200-072) for 10 min at room temperature, and stopped with 500 µL of Dulbecco's modified Eagle's medium (DMEM) + glutamax (Life Technologies, 31966047) containing 10% fetal calf serum (FCS) (Sigma, F7524). Serial 10-fold dilutions from the supernatant and cell suspensions were plated in their specific plate media and grown at 37°C for 3 to 5 days under anaerobic conditions. Colonies were counted to calculate CFU/mL. The following formula was used to calculate the percentage of adherent bacteria:

% attached bacteria = (CFU cell suspension / (CFU cell suspension + CFU supernatant)) * 100

### Peptide nucleotide acid (PNA) probe in-silico design

A collection of 16S ribosomal RNA (rRNA) sequences of different *Prevotella* spp. and related bacteria (*Porphyromonas* spp. and *Bacteroides* spp.) were collected from the Arb-Silva database (https://www.arb-silva.de/search/) and the Ribosomal Database Project (RDP-II, http://rdp.cme.msu.edu/). Sequences were aligned using the ClustalW Omega tool (https://www.genome.jp/tools-bin/clustalw). Probe regions were selected based on sequence homology between strains of interest and mismatches with other strains. Criteria for selection of the most suitable PNA probe included: Gibbs free energy between -13 kcal/mol and 20 kcal/mol, lack of self-complementary structures, length between 15 and 20 nucleotides, melting temperatures (Tm) values between 60 and 80°C, and GC content between 40 and 60%. The theoretical sensitivity and specificity of the probes were determined, and the probes were tested using the TestProbe tool from Arb-Silva (https://www.arb-silva.de/search/testprobe/). The sequences with the best theoretical results were ordered at Integrated DNA Technologies (IDT) with a double 8-amino-3,6-dioxaoctanoic acid (AEEA) linker and a Cy3 resulting in PT-Cy3 and PB-Cy3 PNA/FISH probes.

**Fluorescence In Situ Hybridization (FISH) and confocal microscopy**

VK2/E6E7 and End1/E6E7 cells were seeded on glass slides in a 24-well plate at 70% confluency and grown for 4 days until full confluency. The cells were then transferred to the anaerobic chamber, washed twice with anaerobic DPBS and infected with *G. vaginalis, P. timonensis, P. bivia*, or *L. crispatus* in KSF-medium at a MOI of 50 for 18 h at 37°C in anaerobic conditions. Cells were washed 3 times with cold DPBS with $Mg^{2+}$ and $Ca^{2+}$ (Sigma-Aldrich, D8662) and stained with 10 µg/mL Wheat Germ Agglutinin-663 (WGA-633, Invitrogen, W21404), a lectin that stains the epithelial surface, in cold DPBS for 10 min on ice. Cells were then washed 3 times with cold DPBS and fixed by immersing the slides in methanol for 20 min, followed by 4% cold paraformaldehyde in PBS (VWR, J19943) for 10 min at room temperature (RT) and 50% ethanol for 10 min, all at room temperature. Glass slides were stained by placing them upside down in a 30 µL droplet hybridization buffer containing 0.9 M NaCl, 20 mM Tris (pH 7.5), 0.1 % SDS, and 20 % formamide with 1000 nM EUB338-AF488 probe, 600 nM Gard162-AF488 probe, 200 nM PT-Cy3, or 200 nM PB-Cy3 probe. After 2 h incubation at 50°C in a humidity chamber, the slides were washed by immersing in 1mL hybridization buffer for 15 min at RT and twice with 1 mL DPBS + 0.2 % BSA (Sigma, A7030) for 5 min. Then, slides were stained with DAPI at 1:1000 (Invitrogen, D21490) for 10 min and washed 3 times with DPBS for 15 min in total. The slides were shortly immersed in MilliQ water, and embedded in Prolong diamond mounting solution (Invitrogen, P36990). Imaging was performed within 24 h on a Leica SPE-II confocal microscope in combination with Leica LAS AF software. Image analysis was performed using Fiji/ImageJ.

**Lectin staining and confocal microscopy**

For infection experiments and *O*-glycan analysis, epithelial cells were grown on coverslips in 24-well plates and infected with bacteria as described under the FISH protocol. Cells were treated with 200 U/mL of α2,3,6,8,9 neuraminidase A (NEB Bioke, P0722L) and 0.6 U of α1,2,3,4,6-L- fucosidase (Megazyme, E-FUCHS) for 3 h as positive controls for sialidase and fucosidase activity. Monolayers were washed twice with DPBS and fixed with 4% PFA for 30 min at RT. The fixation was stopped by incubation with 50 mM $NH_4Cl$ in PBS for 10 minutes. Cells were washed twice with DPBS before they were incubated with biotinylated lectins Sambucus Nigra (SNA) (Vector Laboratories, B-1305-2) at 1:200 , Maackia Amurensis Lectin II (MAL-II) (Vector Laboratories, B-1265-1) at 1:100 , and Ulex Europaeus Agglutinin I (UEA-1) (Vector Laboratories, B-1065) at 1:100 in 0.2% BSA in DPBS for 1h at RT. Coverslips were washed 3 times with 0.2% BSA/PBS followed by incubation with Streptavidin-488 (ThermoFisher, A6374) at 1:100 and DAPI at 1:1000 for 1h at RT. Coverslips were washed 3 times with DPBS, once with MilliQ (to remove salts that could interfere with microscopy visualization), and embedded in Prolong diamond mounting solution (Invitrogen, P36990). Images were collected on a Leica SPE-II confocal microscope in combination with Leica LAS AF software unless stated otherwise.

**Cytotoxicity assays**

VK2/E6E7 and End1/E6E7 cells were seeded at 70% in 96-well plates and grown until full confluency and transferred to the anaerobic chamber as described above. Overnight cultures of *P. timonensis*, *P. bivia*, *G. vaginalis,* and *L. crispatus* were washed twice by adding DPBS and centrifugation and resuspended in KSF medium. To determine cytotoxic effects of the different bacteria, cells were infected with bacteria at a MOI of 10 or 100 and incubated for 18 h under anaerobic conditions. The supernatant was transferred to a new 96-well plate and centrifuged at 3400 g for 15 min to remove

remaining bacteria. The presence of released LDH in the supernatant was assessed using the Cytotox 96 Non-Radioactive Cytotoxicity Assay (Promega, G1780). 50 µl Cytotox reagent was added to 50 µl supernatant and incubated for 30 min at room temperature. The reaction was stopped by adding 50 µl of stop solution and absorbance was measured at 492 nm on the FLUOstar Omega microplate reader (BMG Labtech). LDH levels were also determined in a lysate of uninfected cells to determine maximum levels of LDH release. KSF medium background absorbance was subtracted from the measurements and values were averaged from triplicates. The percentage of cytotoxicity was calculated as follows:
% Cytotoxicity = (Experimental LDH release / Maximum LDH release) * 100

**Reverse transcription quantitative polymerase chain reaction (RT-qPCR)**

VK2/E6E7 and End1/E6E7 cells were seeded at 10% in a 12-well plate and grown until 70% confluency and transferred to the anaerobic chamber as described above. Overnight cultures of *P. timonensis, P. bivia, G. vaginalis,* and *L. crispatus* were washed twice with DPBS and resuspended in KSF medium. Cells were infected with bacteria at a MOI of 10 and incubated for 18 h at 37°C in anaerobic conditions. As a positive control, 10 µg/mL Pam3CSK4 (InvivoGen, tlrl-pms) was added to the cells. The next day, cells were washed twice with cold DPBS and RNA was extracted using the RNeasy Mini Kit (Qiagen, 74104). Briefly, cells were lysed in lysis buffer with 10 µL/mL β-mercaptoethanol (Sigma-Aldrich, M3148) and passed 10 times through a 26-gauge needle. After adding 70% ethanol (1:1 ratio), the lysate was transferred to the spin column and RNA was extracted following Qiagen's protocol. RNA quantity and quality was assessed by the spectrophotometer BioDrop µLite (Biochrom) and gel electrophoresis. 2 µg of RNA was treated with DNAse I (ThermoScientific, EN0521), diluted to 10 ng/µL in DEPC-treated water and used for the RT-PCR reaction. The reaction mixture contained 50 ng RNA, 0.3 mM final concentration forward and reverse primers (Table 3), reverse transcriptase (Eurogentec, RT-RTCK-03, 5 U/µl), RNase inhibitor (1 U/µl), 1x No Rox SYBR mastermix blue dTTP (Takyon, UF-NSMT-B0701), supplied with DEPC-treated water to a final volume of 20 µl. A negative control reaction was performed using MilliQ water instead of RNA. RT-qPCR reactions were performed using a LightCycler 480 II system (Roche). Synthesis of cDNA was performed at 48 °C for 30 min, followed by incubation at 95 °C for 5 min to activate the enzyme. Amplification was carried out in 45 cycles of two steps (95 °C for 5 sec and 60 °C for 30 sec). Melting curves were created as follows: 95 °C for 5 sec, 65 °C for 1 min, and a final step at 97 °C. Results were analysed using the LightCycler 480 software. All cycle thresholds were averaged from triplicate reactions and normalized to the housekeeping gene TMEM222. Fold changes were calculated using the delta-delta Ct method.

**Mucin purification**

Mucins for bacterial growth assays were purified from commercially available porcine gastric mucins (PGM, Sigma-Aldrich, M2378). 10 g of PGM was dissolved at 0.5% w/v in 274 mM NaCL, 5.4 mM KCl, 20 mM $Na_2HPO_4$, and 3.6 mM $KH_2PO_4$ (pH 7.8) supplemented with 0.02% v/v toluene. This solution was stirred for 24 h at 4°C during which the pH was adjusted to 7.2 with 2M NaOH after the first hour. The mucin solution was centrifuged at 10,000 x g for 20 min at 4°C, and ice-cold 100% ethanol was added to the supernatant at a final concentration of 60% v/v to precipitate the mucin fraction (without centrifugation). From then on, all steps were performed on ice. After precipitation, the supernatant was discarded, and the mucin precipitate was taken up in 0.1 M NaCl and ice-cold 100% ethanol was added at a final concentration of 60% v/v for a second round of precipitation (without centrifugation). The mucin precipitate was washed with ice-cold 100% ethanol and was shortly incubated to ensure precipitation. The final pellet was dissolved in 180 mL distilled water. The mucin solution was dialyzed against 6 L of distilled water for 24 h at 4°C using snakeskin 10K MWCO (ThermoFisher). The dialyzed mucin solution was frozen horizontally at -80°C in 50 mL tubes to ensure optimal sublimation rates. The frozen mucin solution was lyophilized using a freeze-dryer until all water was evaporated and stored at -80°C until use.

#### Bacterial whole genome sequencing and data processing

Strains were grown on plates of their respective medium and DNA was isolated using the NGS DNeasy UltraClean Microbial Kit (Qiagen, 10196-4) following the manufacturer's instructions. DNA sequencing libraries were prepared using the SQK-RBK110.96 rapid barcoding kit. Nanopore sequencing was conducted using a MinION with R9.4.1 flowcells (Oxford Nanopore Technologies). Basecalling, barcode splitting, and barcode removal were performed using MinKNOW v5.4.3 with the "Super accurate" model. Reads were assembled to contigs with Flye v2.9 (*2*). Assemblies were polished using Medaka 1.4.3 (https://github.com/nanoporetech/medaka) and Homopolish 0.3.4 (*3*). Genomes were annotated using Bakta (*4*).

#### CAZyme analysis

Predicted bacterial protein sequences were used to analyze the presence of carbohydrate-active enzymes using the CAZy database and dbCAN3 meta server (https://bcb.unl.edu/dbCAN2/blast.php). CAZymes identified with at least two out of three tools (HMMER: dbCAN, DIAMOND: CAZy, and HMMER: dbCAN_sub) were considered for further analysis. Signal IP6 server (https://services.healthtech.dtu.dk/services/SignalP-6.0/) was used to predict the presence of signal peptides in the identified proteins.

#### Enzymatic activity assays

Bacteria were grown overnight or until they reached stationary phase and diluted to OD600 = 1 in a final volume of 10.5 mL. The tubes were centrifuged at 8,000 x g for 15 min and the supernatant was concentrated 7.5 times using a 10 kDa spin-X UF 20 mL filter tubes (Corning, 431488). The bacterial pellet was washed once in DPBS and resuspension in 1.4 mL DPBS (resulting in a 7.5 concentration similar to the supernatant fraction). Both the pellet and the supernatant fractions were frozen at -20°C until use. Fucosidase activity was measured by adding 100 µM 4-Methylumbelliferyl α-L-fucopyranoside substrate (Sigma-Aldrich, M8527) to 50 µL of pellet or supernatant samples and incubated for 1 h at 37°C in the dark. Fluorescence was measured at 340 nm (excitation) and 490 nm (emission), and a gain of 1124 (FLUOstar Omega). As a negative control, 100 µM of the fucosidase inhibitor L-fuconojirimycin (FNJ) (Carbosynth, CAS 99212-30-3) was added to the samples and incubated for 30 mins at 37°C in the dark before adding the substrate. Sialidase activity was measured by adding 100 µM 4-Methylumbelliferyl N-acetyl-a-D-neuraminic acid sodium salt (Sigma, M8639) as a substrate. As a negative control, samples were heat-inactivated for 10 min at 95°C. The plates were incubated for 30 mins at 37°C in the dark and fluorescence was measured at 340 nm (excitation) and 490 nm (emission) with a gain of 1124 (CLARIOstar Plus). For all assays, measured values for DPBS or the relevant media were subtracted from the sample measurements.

#### Cloning of *P. timonensis* sialidase genes

Full-length *P. timonensis* sialidase genes were amplified using Q5 polymerase (New England Biolabs, M0492S) from genomic DNA purifications (DNeasy kit, Qiagen, 12224-50) using primer pairs depicted in Table 4. The resulting gene products were assembled into pET28a expression vector (Novagen, 69864) using Gibson assembly (New England Biolabs, E2611S) and transformed into DH5α *Escherichia coli* chemically competent cells. The resulting plasmids pET28-6xHis-NanH1 and pET28-6xHis-NanH2 were transformed into *E. coli* BL21 (DE3) (New England Biolabs, C2530H). Transformed bacteria were grown in Luria Broth (LB) (Research Products International, L24060-2000) with 50 µg/mL kanamycin sulfate (VWR, 408-EU-25G). Plasmids were purified with Plasmid Midiprep (Zymo Research, D4201) and insert sequences were confirmed by DNA sequencing (Genewiz).

Full sequence of PtNanH1:
MKIFRLISFLWLLSTGLVVQASNNQHHQQLLFETDSVNKIPYRIPAIAQCRNGNLIAVSDFRYC
GSDIGYGAVDLVYRISKDYGITWSPISKLADGHGDENRIQWDYAFGDCGLVANRTNSEVLAV
CVAGKTVYFQGKRNNPNRVAVFRSNDNGKTWDKGHEITEQIYRLFDGRKQGPIQSLFFTSGR
IHQSRYVKVGKYFRLYSGLCTLSGNFIVYSDDFGHQWKVLGNIDESPCKDGDEVKCEELPDG
SLILSSRTEGRMFNIFTFTNIKKGEGHWDSRQTASDMAHIKNQCNGEILVLPVIRKSDGQETYL
ALQSVPFGPQRTNVGIFYKEIGKEHLSALEFASHWKRGMQVHHGPSSYSTMIAQKDGRIGFL
CEVGQKNNHISYQSLDVEEITNNEFVLSKKRFKKGY*

Full sequence of PtNanH2:
MNLKSLRLLWTFALLLALGWSTAHAADKVIRIQKSTGTWTKSNPAKTWAAQWTSNDVDPM
LTLTCAANNMAYYDGNEIKLFTGNGSVKFSADYTLAVIPGYEIVSYEFSFSSEKAGQKIVVTP
QGATELSSDDPTQWQAVQVTDIHAATATFNVKHATQTAAGFARLKNVLVTVAKTATPANF
HPLYITRPNEHPYRIPALACLKDGKLLAFSDYRPGGSDIGYGEVDIQLRTSTDHGATWTDART
IANGKGTGSGLDYGFGDAAVVADRESNEVLMMCVAGHVPYQTANYQAGNPLPMVRYLST
DGGETWGNYTDVTAAVYGLFDGSKDGAVKSMFAGSGKLCQSSIVKHGTHYRVYMPVCAR
DGGNRVLYSDDFGATWKVLGGKDARPAPGGDEPKCEELPDGRVLLSSRTNGGRLFNIYTYT
NHETGEGSWGEVAKSDANNRGTAAQSNSCNGEIMILPVKRNSDGKQMFLALQSVPFGPNRA
NVGIYFKALESLNDYFTPKAFASNWTGKKQISQMGSAYSTMVWQKDDKVGFFYEESTYGA
DYTNVYKALTLEEITDNAFSYDATVKRPTEDANAIQYSDLTKVTQLLGHKGVGFPLADAESR
ELLAKIYDAPANYTKAQLEKAVQQFVEETNVEKPQNGLLYLVKFVGKDGKTYMVDYADGK
LSAKPITEGQAPERSAGFKAHVLATGKVAFETMDGKFLCYPTQLPAPQDLTGYHVGGVTDEL
DEADNGLDLQKAIASDKVESTEALDRFGKFYICSKRGNRTDNQQEALGFWTLNTQANAFNN
ADVPFLNNQLTSLVVVERMVLPLGKQVTTLAEVDPNKAYALYNEHFTTYAVKKEGQTNVW
VQGMVGDAGHSLANGDFAQPYNPSSAFGAWQLMKNDQGQWLLYNIGAKQYARTPSNQSG
TGPCTFVDEALPINVTELPNGGFAFNTGNNAQQFFCAAPQLADSPINVWTSSDSGSRWLLVE
NPNVQVDVPNSIQQTVAAKANRKTGIYNLKGQRINTDKLHQLPAGIYIVDGKKVVVK*

**Recombinant expression and purification of *P. timonensis* sialidases**

For expression of the *P. timonensis* sialidases, BL21 (DE3) containing pET28-6xHis-NanH1 or pET28-6xHis-NanH2 were grown at 37 °C with shaking (180 rpm) in 1 L of LB containing 50 µg/mL kanamycin. When the culture reached an $OD_{600}$ between 0.4-0.8, protein expression was induced by adding Isopropyl β-D-thiogalactoside (IPTG) (Teknova, I3325) to a final concentration of 250 µM. The cultures were incubated overnight at 16 °C with shaking (180 rpm). Cells were then harvested by centrifugation at 6000 x g for 10 minutes at 4 °C. Cell pellets were resuspended in Buffer A (20 mM Tris, 500 mM NaCl, 10 mM $MgSO_4$, 1 mM $CaCl_2$, pH 7.5) supplemented with 0.1 mg/mL DNAse (Sigma, DN25-1G), 0.5 mg/mL Lysozyme (L6876-10G), and Pierce Protease Inhibitor Tablets (Thermo Scientific, A32965). Cells were lysed using a cell disrupter (Emulsiflex-C3, Avestin, C315661) by passing four times at 15,000 psi. Lysates were clarified by centrifugation at 9,000 rpm for 45 minutes at 4 °C. The clarified lysate was transferred to a 15 mL column containing 3 mL Ni-NTA Agarose affinity resin (Qiagen, 1018236) and washed with 12 mL Buffer A. The column was washed using Buffer A with a gradient of 10 mM, 75 mM, and 100 mM imidazole. Protein was eluted with Buffer A containing 250 mM imidazole. Fractions were analyzed by SDS-PAGE (Biorad, 4561086) and stained with Instantblue Coomassie protein stain (Abcam, ab119211). The protein-containing fractions were pooled and concentrated in an Amicon spin concentrator of 10-30kDa cutoff (Millipore, UCF901024 and UCF903024). Size exclusion chromatography was using a ÄKTA pure (Cytiva). 2 mL of enzyme solution was loaded into a HiLoad 16/600 Superdex 200 pg size exclusion column (Cytiva, 28989335) pre-equilibrated in Buffer A. Aliquots were frozen in liquid nitrogen and stored at –80 °C. Protein concentrations were determined by UV-Vis (Agilent, G9864A) and the extinction coefficient was calculated using the following online tool: https://www.novoprolabs.com/tools/protein-extinction-coefficient-calculation. To verify the recombinant purified sialidases, SDS-PAGE gels (Biorad, 4561086) were run and stained with Instant Blue Coomassie protein stain (Abcam, ab119211) (Figure S6 of Pelayo *et al*., accompanying paper).

**Statistical analysis**

Statistical analysis was performed using Graph Pad Prism 7 software. The Kolmogorov-Smirnov test was used to assess normality of the data, and log transformation was used when the data was not normally distributed. The bacterial adhesion capacity, and LDH release data were analyzed using the Dunnett correction (by comparing each bacterium to the control *L. crispatus*). Data from RT-qPCR and bacterial growth experiments were assessed by the two-way ANOVA (analysis of variance) test with Dunnett's correction (comparison of each bacterium to the control *L. crispatus*). Statistical differences in data of enzymatic activity assays were analyzed using one-way ANOVA with Tukey's HSD *post hoc* test. Microscopy images quantification was done using one-way ANOVA with Dunnett's correction and compared to the untreated cells (control). All graphs depict the mean and standard error of the mean (SEM) of at least three independent experiments. A *p*-value of <0.05 was considered significant. * $p<0.05$; ** $p<0.01$; *** $p<0.001$. **** $p<0.0001$.

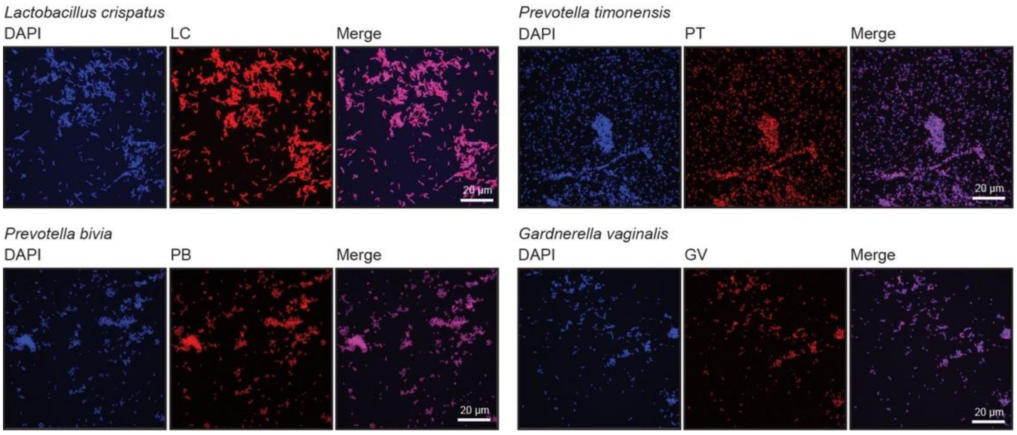


**Figure S1. Confirmation of FISH probe staining of vaginal bacteria**

Confocal microscopy images of vaginal bacteria attached to glass slides stained with peptide nucleic acid (PNA) probes listed in Table 2. For each bacterium, the corresponding PNA signal is shown in red while DAPI is shown in blue. White scale bars represent 20 µM.
